# A fibril-inducing support-bath enables embedded 3D printing of aligned collagen-rich constructs

**DOI:** 10.64898/2026.08.30.748057

**Authors:** Giovanni Gonnella, Rosario Milazzo, Rory Gibney, Daniel J. Kelly

**Affiliations:** Trinity Centre for Biomedical Engineering, Trinity Biomedical Sciences Institute, Trinity College Dublin, Dublin, Ireland; Department of Mechanical, Manufacturing and Biomedical Engineering, School of Engineering, Trinity College Dublin, Dublin, Ireland; Department of Anatomy and Regenerative Medicine, Royal College of Surgeons in Ireland, Dublin, Ireland; Advanced Materials and Bioengineering Research Centre (AMBER), Royal College of Surgeons in Ireland and Trinity College Dublin, Dublin, Ireland

## Abstract

Embedded extrusion printing can process collagen-rich bioinks, but their low viscosity and slow fibrillogenesis compromise print fidelity and post-deposition stability. Here, we developed a collagen fibril-inducing support bath (FIB) that combines mechanical support for embedded printing with biochemical induction of collagen assembly. Microfibrillated or nanofibrillated cellulose was incorporated into a fibril-inducing buffer, and formulations were screened at 37 °C for rheological behaviour and optical transparency. The selected FIB was evaluated by printing 1% and 5% (w/v) articular cartilage-derived extracellular matrix (ECM) inks at 10–20 mm s□¹ and compared with a cellulose-only control bath. FIB exhibited yield-stress, shear-thinning and rapid recovery behaviour that supported reproducible filament deposition. Unlike the control bath, FIB enabled intact construct retrieval following stabilisation and promoted the formation of fibrillar collagen within the printed strands. Scanning electron microscopy revealed D-banded collagen fibrils preferentially oriented along the deposition direction, with dominant orientation peaks within ±10–15°. The platform supported the fabrication of 15 × 15 × 1.5 mm sheets and 6 × 6 × 6 mm scaffolds whose macroscopic dimensions were retained after processing. Constructs produced from 5% ECM inks exhibited approximately fourfold higher ramp and relaxation moduli than those produced from 1% ECM inks. Extracts from both formulations caused no detectable reduction in cell metabolic activity after 24 or 72 h. Mesenchymal stem/stromal cells (MSCs) seeded onto printed sheets became markedly elongated and aligned by day 3, with approximately 80% of cells having an aspect ratio exceeding 1.5, significantly greater than cells seeded onto casted ECM controls, with a mean deviation of ∼9° from the filament print direction. These findings establish FIB as a bioactive support bath that couples embedded printability with collagen fibrillogenesis, enabling recoverable collagen-rich constructs with aligned fibrillar architecture that directs early cellular organisation.

## 1. Introduction

Collagen fibre orientation and spatial patterning regulate cell behaviour and are integral to the mechanical function of diverse tissues and organs [1–4]. Collagen exhibits tissue-specific architectures, for example, a lamellar organisation in cornea [5,6], helical fibre patterns in arteries [7,8], and arcade-like collagen fibres in articular cartilage [9,10], which collectively highlight how collagen fibre organisation and alignment underpin tissue function.

Collagen hydrogels are widely used in tissue engineering as they are biocompatible and support cell survival, growth, and differentiation [11,12]. Gelation occurs *via* collagen fibrillation, in which soluble monomers self-assemble into fibrils and higher-order networks [13–15]. Fibrillation of collagen is governed by pH, ionic strength, temperature, collagen concentration, and molecular crowding, which together regulate assembly kinetics and the resulting network structure and mechanics [15–18]. Because collagen microstructure and alignment influence cell migration, proliferation, differentiation and matrix deposition, strategies to generate aligned and/or spatially organised collagen-based biomaterials have been widely explored [19–23]. However, many approaches are limited to planar formats or restricted geometries and do not readily enable the biofabrication of complex 3D constructs at relevant dimensions [19,24].

Extrusion-based 3D (bio)printing offers scalable fabrication of scaffolds and cell-laden constructs with spatial control over geometry and material placement [25–29]. Furthermore, the alignment of fibrillar components within (bio)inks can be partially controlled by modulating the shear forces they experience during extrusion-based deposition [5,30]. The printing of collagen is challenging, due in part to its relatively low viscosity and poor mechanical properties prior to (or during early stages of) collagen fibrillation [31,32]. As a result, filaments can spread and/or collapse after extrusion, leading to poor shape fidelity and limiting the fabrication of stable, complex architectures [28,31].

Embedded printing of collagen within yield-stress, shear-thinning support-baths mitigates collapse and spreading by providing mechanical confinement during deposition [33–37]. However, many support-baths are optimised for short-term geometric support and subsequent removal rather than actively promoting collagen fibril and fibril organisation at physiological temperature [38,39]. Gelatin microparticle baths are thermoresponsive and often require temperature control for printing and/or removal, which can constrain workflows that require extended incubation at 37 °C for collagen-rich inks to develop stable fibrillar networks [34,40]. Alternative polymeric baths (e.g., xanthan-based systems [41,42]) can provide suitable rheology for embedded printing, but may require chemical modification or post- processing that complicates construct retrieval, particularly for porous architectures [34,35,42]. Cellulose-based fibrillar suspensions are attractive due to their biocompatibility and tunable rheology [43,44], yet systematic relationships between formulation, optical transparency, and collagen print fidelity have yet to be defined. Collectively, these limitations motivate a support-bath that (i) stabilises low-viscosity, collagen-rich inks during printing, (ii) supports collagen fibril and alignment, and (iii) enables straightforward construct recovery.

Here, we repurposed a previously reported collagen fibril–inducing buffer [45] as the basis for a collagen fibril-inducing support-bath (FIB), tuning yield stress and recovery using microfibrillated or nanofibrillated cellulose and evaluating its suitability for embedded printing of collagen-rich inks. (Note that throughout this manuscript, the term filament refers to the macroscale strand deposited during extrusion printing, whereas collagen fibril refers to the nanoscale collagen structures formed through fibrillation within the printed filament. The term collagen fibre is reserved for the higher-order collagen organisation found in native tissues). We quantified bath rheology at physiological temperature and optical transparency across formulations, then systematically assessed the printing of low- and high-concentration extracellular matrix (ECM)-based inks at different printing speeds at a fixed flow rate, benchmarking against a cellulose-only control bath. Finally, we attempted to leverage this support-bath platform to fabricate multilayer architectures, including large sheets (15 x 15 x approximately 1.5 mm) and cubic scaffolds (6 x 6 x 6 mm) with programmed filament spacing. Collagen alignment, post-processing stability and compressive mechanical properties were subsequently assessed. Eluate-based MTT testing and phalloidin staining of mesenchymal stem/stromal cells (MSCs) were used to evaluate construct cytocompatibility and the ability of the printed filaments to guide early cellular elongation and orientation. Together, this work establishes a support-bath platform for embedded 3D printing of collagen-rich inks that promotes fibrillation, fibril alignment, programmable filament organisation, improved post-deposition stability, and enables straightforward construct retrieval while supporting early cell organisation.

## 2. Methods

### 2.1. Preparation of the support-bath

To prepare the fibril-inducing support-bath, a previously developed fibril-inducing support buffer [45] was modified by adding microfibrillated cellulose (MFC, Nanografi, Turkey) or nanofibrillated cellulose (NFC, Nanografi, Turkey) at varying concentrations to increase viscosity. First, a 500 mM phosphate buffer (PB) was prepared by dissolving sodium phosphate dibasic heptahydrate (377 mM, Sigma) and sodium phosphate monobasic monohydrate (123 mM, Sigma) in ultrapure water. The fibril-inducing buffer was then formulated by combining an aliquot of the 500 mM PB stock with ultrapure water to obtain a final phosphate concentration of 118 mM, followed by addition of 20% (w/v) polyethylene glycol (PEG; MW 8000; Sigma). To generate cellulose-thickened fibril-inducing support-bath formulations, MFC or NFC was dispersed into the fibril-inducing buffer to final concentrations of 2% (w/v) MFC, 10% (w/v) NFC, or 15% (w/v) NFC. The final solution was stirred on a magnetic stir plate overnight at room temperature to ensure complete homogenisation. Final pH was then adjusted to 7.4 prior to utilisation.

### 2.2. Rheological and optical characterisation of the support-bath

Rheological characterisation were conducted on an MCR 102 rheometer (Anton-Paar, Hertford Herts, UK) equipped with a Peltier element for temperature control. A parallel-plate geometry with a width of 25 mm (PP25) was used in all tests. The viscosity as a function of shear rate (0.1 – 1000 s^−1^) was investigated at a constant temperature of 37 °C. The ability of the bath to perform recovery after shear was assessed by sequential cycles of low and high shear strains, evaluating the relationship between G’ (storage modulus) and G” (loss modulus). The amplitude sweep was carried out at a fixed oscillation frequency of 1 Hz over 0.01 – 1000 % shear strain at 37 °C. As a control, 2% nanofibrillated cellulose dispersed in ultrapure water was used. n = 4 samples per groups were used during this test.

To assess optical properties of the support-bath, 100 µL of solution was initially transferred into a clear flat-bottom 96 well-plate (Sigma). 1X Phosphate buffer saline (PBS) solution was used as a blank for this experiment. Optical transparency/opacity was quantified by UV–Vis spectrophotometry (BioTek plate reader, Agilent Technologies, USA) by measuring absorbance (and calculating % transmittance) across the visible range (350–700 nm), as commonly reported for hydrogel and nanocellulose-based materials [46,47].

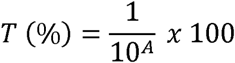

n = 4 samples per groups were used during this assay.

### 2.3. Articular cartilage ECM extraction

Articular cartilage (AC) was harvested from the condyles of 6-month-old pigs following a previously established protocol [48,49]. Briefly, articular cartilage was removed from the tibia and femur using 6 mm biopsy punches and finely diced into small pieces (3–4 mm) using a commercial knife. The tissue was weighed, transferred into 50 mL conical tubes, and pre-treated with 0.2 M NaOH for 24 h at 4 °C to remove non-specific proteins and sulfated glycosaminoglycans (sGAGs) [50,51]. After pre-treatment, the NaOH solution was removed, and the cartilage was thoroughly washed with ultrapure water. The extracellular matrix (ECM) was then enzymatically extracted using a solution of 1500 U/mL of pepsin (Sigma) in 0.5 M acetic acid (HAc, Sigma), under slow rotation at 4 °C for 24 h. The tubes were centrifuged at 2500 x g for 1 hour at 4 °C to remove non-solubilised material, with the pellet discarded and the supernatant retained. The supernatant, containing solubilised ECM, was transferred to new conical tubes, and type II collagen fibers were salted out by adding 5 M sodium chloride (NaCl, Sigma) to a final concentration of 0.9 M NaCl overnight at 4 °C [52]. The tubes were centrifuged at 2500 g for 1 hour at 4 °C, and the supernatant was discarded. The collagen pellet was resuspended overnight in 0.5 M HAc at 4 °C. The salt precipitation step was repeated once more to increase ECM purity. The solubilised ECM was then transferred to 6–8 kDa dialysis membranes (Spectrum Labs) and dialyzed against ultrapure water for 48 h, with the solution changed every 24 h. Finally, the ECM was transferred into petri dishes and lyophilized using a Labconco (FreeZone Triad, Labconco, KC, USA) freeze-drier. The lyophilized ECM was stored at -20 °C until further use.

### 2.4. Preparation of AC-ECM inks for 3D printing

AC-ECM inks were fabricated following a previously established protocol[49]. Briefly, lyophilized ECM was placed into 5 mL tubes and resuspended in high-glucose Dulbecco’s Modified Eagle Medium (DMEM, Gibco) to achieve final concentrations of 10 mg/mL (1%) and 50 mg/mL (5%). To allow ECM solubilisation, 0.5 M acetic acid was added to a final concentration of 0.02 M. Tubes were rotated at 2 rpm for 2 days at 4 °C to allow complete solubilisation. After solubilisation, the solutions were neutralized with small volumes of 0.1 M NaOH and chemically crosslinked with 100 mM glyoxal (Sigma) to a final concentration of 5 mM. Crosslinking was performed by incubating the solutions at 37 °C for 30 minutes before loading them into 3 mL syringes for 3D printing.

### 2.5. Rheological analysis of AC-ECM inks

Before proceeding to 3D printing, the rheological properties of the two inks were assessed following the same procedure as described in *section 2.2*. Briefly, the viscosity as a function of shear rate (0.1 – 1000 s^−1^) was investigated at a constant temperature of 37 °C. The ability of the inks to perform self-recovery was assessed by sequential cycles of low and high shear strains, evaluating the relationship between G’ (storage modulus) and G” (loss modulus). The amplitude sweep was carried out at a fixed oscillation frequency of 1 Hz over 0.01 – 1000 % shear strain at 37 °C.

### 2.6. 3D printing of AC-ECM inks into fibril-inducing support bath

All printing experiments were performed using a Cellink BIO X6 bioprinter (BICO, Sweden) equipped with a syringe pump extrusion system. A 3 mL syringe fitted with a 27 G needle (inner width = 210 μm) was used, and the printing bed was maintained at 37 °C. Volumetric flow rates were fixed at 3.5 µL/s for the 5% AC-ECM ink and 3.0 µL/s for the 1% AC-ECM ink.

Printing fidelity was assessed in a stepwise manner, following previous methods [5,53–55]. Briefly, to quantify filament spreading, straight filaments were printed at translation speeds of 10, 15, and 20 mm/s for each ink concentration (5% and 1%). For each condition, three independent prints were performed (n = 3), with three parallel filaments printed per replicate. Filaments were imaged immediately after printing using a stereomicroscope (Ash Inspex HD 1080p), and filament width was quantified in ImageJ. For each printed filament, the width was measured at three positions along the filament length (3 filaments × 3 positions) and averaged.

To assess print survivability and post-processing dimensional changes, the single-filament printing experiment was repeated and the printed bath–construct assemblies were frozen at - 80 °C and lyophilised, followed by dehydrothermal (DHT) crosslinking at 115 °C and 2 mbar for 24 h. Constructs were subsequently rehydrated in ultrapure water and re-imaged, and filament width and filament length were quantified post-DHT/rehydration to evaluate dimensional stability and swelling relative to measurements acquired immediately after printing.

To determine printing parameters for porous versus solid architectures, a line-spacing study was performed. Filament-to-filament spacing was varied from 0.20 to 0.80 mm in 0.05 mm increments, with an additional 1.0 mm spacing included as a non-overlapping control. To assess the impact of stacking on strand fusion, patterns were printed either as a single layer (layer height 0.2 mm) or as three layers (three stacked layers at 0.2 mm layer height each). Based on the single-filament optimisation, printing speed was fixed at 15 mm/s for the 5% ink and 20 mm/s for the 1% ink. For each spacing and layer condition, three independent prints were performed and imaged immediately after printing. Fusion behaviour was scored in ImageJ as overlapping, touching, or separated for adjacent filaments, and the minimum spacing required to achieve separated filaments (i.e., the non-fusing threshold) was compared between the single- and three-layer patterns.

### 2.7. Scanning electron microscopy and collagen fibrils alignment determination

Prior to scanning electron microscopy (SEM), printed constructs were dried following previous methods [56,57]. Samples were fixed in paraformaldehyde (4% w/v in PBS) overnight at 4 C, dehydrated through a graded ethanol series (50%, 70%, 90%, 100%), exchanged into hexamethyldisilazane (HMDS), and air-dried. Prior to SEM imaging, samples were sputter-coated with a gold/palladium alloy (Agar Scientific, UK) for 90 seconds at 0.1 mBar using a 108 Auto Sputter Coater (Cressington, UK). SEM images were captured using a Zeiss Ultra Plus (Zeiss, Germany) with an acceleration voltage of 5 kV and a working distance of 5 mm. To quantify the collagen fibril alignment, 5 images per sample were taken (n = 3 samples) and the alignment was quantified via the ImageJ directionality plugin.

### 2.8. 3D printing of representative geometries

Representative geometries were printed in the fibril-inducing bath (FIB) using the printing parameters identified in the filament fidelity studies, *section 2.6*. A cube construct (6 × 6 × 6 mm) was printed using both the 1% and 5% AC-ECM inks with a depth-dependent spacing strategy to program filament spacing as a function of construct height. For the 1% ink, the bottom region was printed with a filament-to-filament spacing of 1.5 mm to generate large pores (3 mm height), the mid-region spacing was reduced to 1.0 mm (2.4 mm height), and the surface region spacing was further reduced to 0.65 mm (0.6 mm height). For the 5% ink, the filament-to-filament spacings were 1.3 mm for the basal region, reduced to 0.85 mm for the mid region and ultimately reduced to 0.55 mm for the surface region. This was done to produce a denser top architecture in which filaments were approaching contact, resembling the superficial zone organisation of articular cartilage [9]. Lastly, a sheet construct (15 × 15 × 1.5 mm) was printed using only the 5% AC-ECM ink as a unidirectional (single-direction) line pattern with a filament-to-filament spacing of 0.55 mm, producing a continuous, densely packed architecture. Samples were crosslinked and retrieved following the same procedure as in *section 2.6* and the fiber orientation was assessed via SEM imaging and using the directionality plugin in ImageJ (5 images per sample were taken and quantified).

### 2.9. Mechanical test

Mechanical properties of the cube constructs (n = 3) were assessed using a Zwick Roell compression tester (Herefordshire, UK) equipped with a 5 N load cell. Constructs were kept in a PBS bath to ensure hydration throughout the test and maintained at room temperature. An initial preload of 0.005 N was applied for 60 seconds, to ensure that both surfaces of the samples were in contact with the compression plates. Force was subsequently zeroed, and samples were subjected to unconfined stress-relaxation compression tests at 10%, 20%, and 30% strains, at a strain rate of 2.3% strain per second. A 30-minute relaxation period was added between each step to fully characterise the viscoelastic behaviour of the constructs. The load versus displacement data were recorded throughout. The stress and strain were calculated dividing the load value by the initial cross-sectional area of each sample, and the displacement value by the initial height, respectively.

### 2.10. Isolation and expansion of caprine mesenchymal stromal cells (MSCs)

Bone marrow-derived mesenchymal stromal cells (MSCs) were obtained following a previously established method [58–60]. Briefly, MSCs were isolated from the femurs of a 4- month-old caprine donor acquired from a local veterinary (Lyon’s farm, University College Dublin) following all relevant guidelines and regulations. The bone marrow was extracted from the femoral shaft and washed with growth medium, which consisted of high-glucose DMEM (Biosciences, Ireland) supplemented with 10% fetal bovine serum (FBS, GIBCO, Biosciences, Ireland) and 2% penicillin (100 U/mL) and streptomycin (100 µg/mL) (Biosciences, Ireland). A homogeneous suspension was prepared by triturating the marrow with a needle. The solution was centrifuged twice at 650 g for 5 minutes, with the supernatant discarded each time. The cell pellet was triturated and filtered through a 40 µm cell sieve. Following colony formation, cells were trypsinized, counted, and re-plated at a density of 5 × 10³ cells/cm² for further passage. All cell expansion was conducted under normoxic conditions in growth medium, with media changes occurring twice weekly. Cells were used at the end of passage 3.

### 2.11. Cytotoxicity of the printed constructs

MTT (3-(4,5-dimethylthiazol’-2-yl)-2,5-Diphenyltetrazolium Bromide) was used to measure cellular metabolic activity [61]. Previously expanded MSCs were used for this first screening of the construct’s extracts. In brief, both 1% and 5% AC-ECM cell-free printed cubes were initially placed in elution for both 24 h and 72 h on a roller shaker in expansion medium comprising of high-glucose DMEM (Biosciences, Ireland) supplemented with 10% fetal bovine serum (FBS, GIBCO, Biosciences, Ireland) and 2% penicillin (100 U/mL) and streptomycin (100 µg/mL) (Biosciences, Ireland). n = 3 samples per group were used. MSCs were seeded at 10^4^ cells/well in two 96-well plates and incubated for 24 h. Cells were subsequently incubated with the eluted media for a further 24 h and 72 h prior to the addition of MTT solution (25 mg of MTT was dissolved in 5 mL, which was diluted to 0.5 mg/mL in ascorbic-free expansion medium). The plates were incubated for 4 h, after which they were gently blotted dry, and 100 μL of dimethyl sulfoxide (DMSO) was added to each well. Plates were positioned on a plate shaker for 5 min before placing them in a microplate reader. The absorbance of the samples was estimated at a wavelength of 570 nm. As controls, cells cultured in expansion media (negative control) and in expansion media with 10% EtOH (positive control) were used.

### 2.12. MSCs morphology and alignment on printed sheets

MSCs were seeded onto printed collagen sheets (n = 3) and cast controls (n = 3) at a density of 1.5 x 10^5^ cells/constructs following previous methods [62,63]. After 1 and 3 days of culture, samples were fixed in 4% paraformaldehyde, permeabilised with 1 % Tween (Sigma) in 1x PBS, and stained with phalloidin and DAPI to visualise F-actin and cell nuclei, respectively. Images were acquired using a confocal microscope (Leica) from 3 randomly selected regions of each construct using identical acquisition settings and averaged for each independently fabricated construct. Maximum-intensity projections were analysed in Fiji. Cell morphology on printed sheets and cast controls was assessed following phalloidin staining of the F-actin cytoskeleton. Individual, non-overlapping cells were manually outlined or segmented from the phalloidin channel, excluding cells intersecting the image boundaries. Cytoskeletal elongation was quantified using the aspect ratio (AR), calculated as the major-axis length divided by the minor-axis length. A mean cell AR between 1.0 and 1.5 would indicate rounded morphology and poor cytoskeleton elongation, while an AR higher than 1.5 would indicate cell elongation on the seeded surface [64–66]. The average cells AR, as well as the percentage of cells displaying an AR above 1.5 was calculated, together with the angle of cytoskeleton elongation. Lastly, the angle of cytoskeleton elongation was compared to the collagen fibre direction to quantify the amount (in percentage) of aligned cells on the printed construct. Cell alignment relative to the printed collagen filaments was quantified in Fiji. An ellipse was fitted to each segmented cell, and the orientation of its major axis was taken as the cell orientation angle. The direction of the corresponding printed filament was defined using a straight reference line. The absolute angular deviation (Δθ) between the cellular major axis and the filament axis was calculated as:

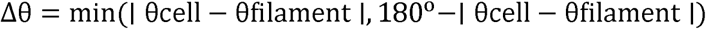

Angular deviations therefore ranged from 0° to 90°, where 0° indicated perfect alignment parallel to the filament and 90° indicated perpendicular orientation [67,68].

### 2.13. Statistical analysis

Results are presented as mean +/- standard deviation. Statistical analyses were performed with two-way analysis of variance (ANOVA) with Tukey’s post hoc multiple comparison test. Statistically significant changes are marked as * p < 0.05, ** p < 0.01, *** p < 0.001, **** p < 0.0001.

## 3. Results

### 3.1. Cellulose-based support-baths exhibit shear-thinning behaviour and rapid recovery at 37°C

To generate cellulose-thickened fibril-inducing support-bath formulations, MFC or NFC was dispersed into the fibril-inducing buffer to final concentrations of 2% (w/v) MFC, 10% (w/v) NFC, or 15% (w/v) NFC (Figure 1a). Rheological analysis revealed that all bath formulations exhibited pronounced shear-thinning behaviour, with viscosity decreasing as shear rate increased (Figure 1b). Under cyclic high- and low-shear conditions, all baths also demonstrated thixotropic recovery, indicating partial restoration of structure after shear-induced breakdown (Figure 1c). Amplitude sweep testing further confirmed predominantly solid-like behaviour at low strain, with G′ exceeding G″ across the linear viscoelastic region, supporting the ability of the baths to provide mechanical confinement during printing (Figure 1d). Additionally, NFC-based support-baths exhibited superior transparency when compared to MFC-based support-baths (Figure 1e, f).

**Figure 1:**
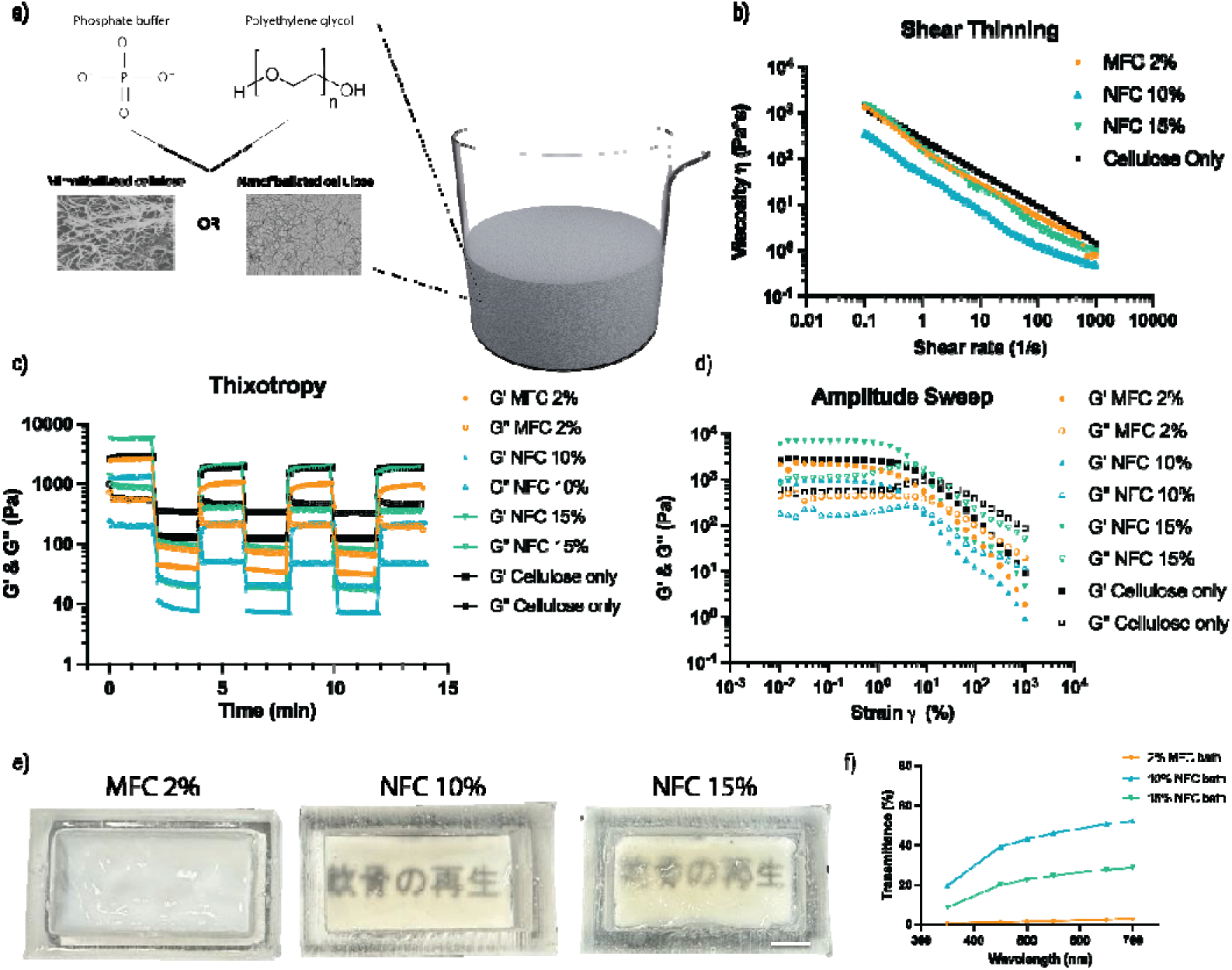
Development and characterisation of NFC- and MFC-based support-baths. Schematic illustration of the support-bath formulation, comprising phosphate buffer, polyethylene glycol, and cellulose reinforcement in the form of MFC or NFC (a). Shear-thinning behaviour of the bath formulations, shown as viscosity as a function of shear rate (b). Thixotropic recovery of the formulations during cyclic step-shear testing, shown by changes in storage modulus (G′) and loss modulus (G″) over time (c). Amplitude sweep analysis showing viscoelastic behaviour of the support-baths across increasing strain (d). Representative images of the different bath formulations demonstrating differences in optical appearance (e). Transmittance spectra of the bath formulations across the visible wavelength range (f). Data are presented as mean ± standard deviation, with statistical significance indicated in the figure. n = 4 samples per group. * p < 0.05, ** p < 0.01, *** p < 0.001, **** p < 0.0001. Scale bar = 1 cm.

### 3.2. Embedded printing supports continuous filament deposition and construct retrieval

Articular cartilage ECM-derived inks were prepared at 1% and 5% (w/v) following decellularisation, pepsin solubilisation, salt precipitation, lyophilisation and reconstitution (Figure 2a). Both inks exhibited shear-thinning behaviour, with viscosity decreasing as shear rate increased (Figure 2b). Across the tested range, the 5% AC-ECM ink had higher viscosity than the 1% AC-ECM ink. During step-shear testing, both formulations showed a decrease in moduli under high shear, followed by recovery under low shear conditions (Figure 2c). In amplitude sweep analysis, G′ and G″ were higher in the 5% ink than in the 1% ink over the low-strain range (Figure 2d).

**Figure 2:**
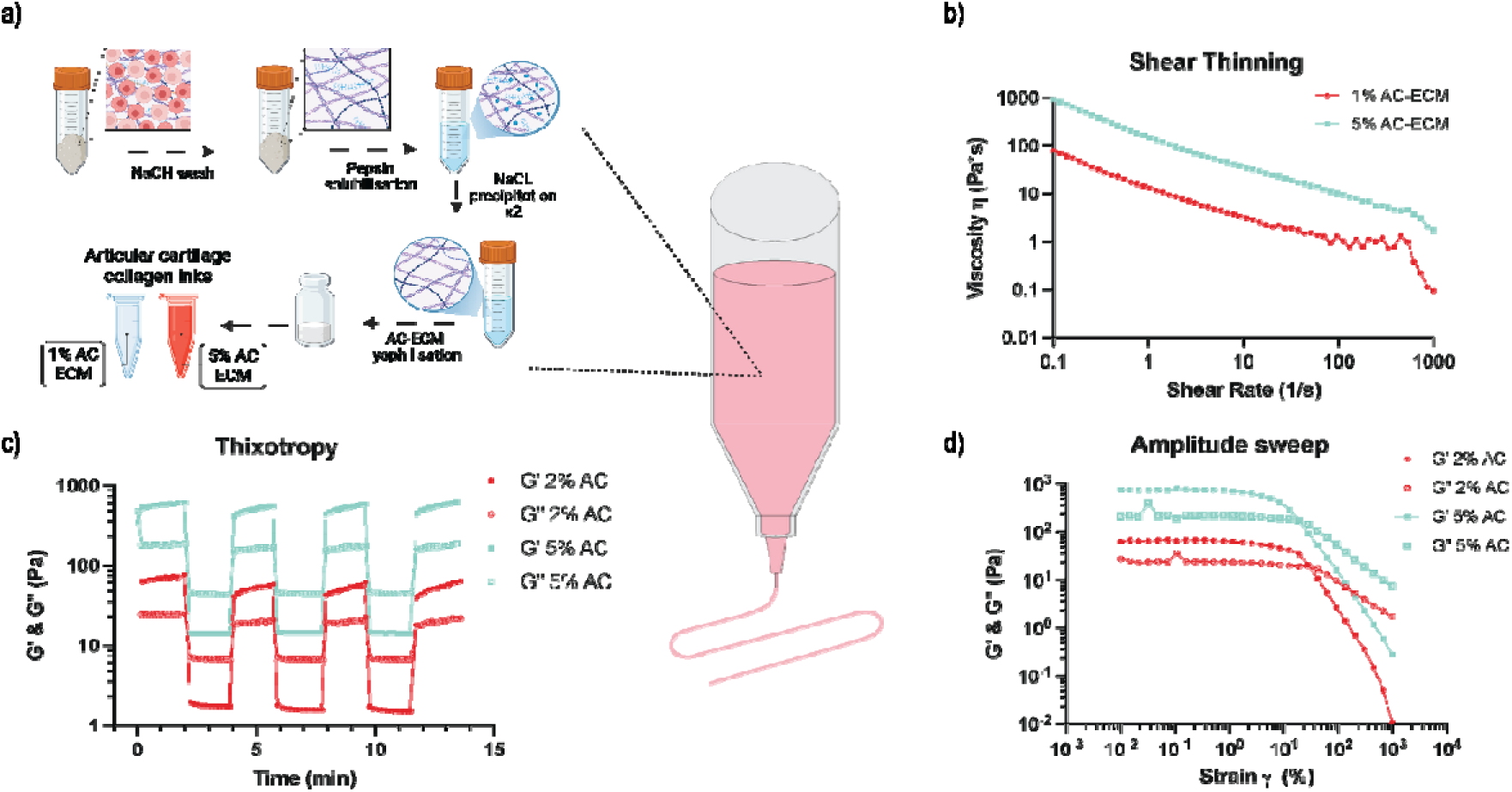
Preparation and rheological characterisation of articular cartilage-derived ECM inks. Schematic illustration of AC-ECM ink preparation, including NaOH washing, pepsin solubilisation, NaCl precipitation, lyophilisation and reconstitution into 1% and 5% AC-ECM inks (a). Viscosity of 1% and 5% AC-ECM inks as a function of shear rate (b). Storage modulus (G′) and loss modulus (G″) during cyclic step-shear testing (c). Storage modulus (G′) and loss modulus (G″) across increasing strain during amplitude sweep analysis (d). Data are presented as mean ± standard deviation. n = 4 samples per group. * p < 0.05, ** p < 0.01, *** p < 0.001, **** p < 0.0001.

Both ECM inks were then embedded-printed in the fibril-inducing bath (FIB) or a cellulose-only control bath (Figure 3a). Immediately after printing, filament width was significantly higher in the FIB bath compared to the cellulose-only bath for both ink concentrations and across all tested print speeds, except for the 1% ink printed at 20 mm/s (Figure 3b). To retrieve the printed samples, the entire support bath containing the constructs was first frozen, freeze-dried and subjected to dehydrothermal treatment (DHT). The dried bath was then immersed in distilled water, causing the bath material to precipitate while the collagen-rich constructs remained buoyant and floated to the surface, enabling their gentle recovery. Post-processing analysis after freeze-drying and dehydrothermal crosslinking treatment was restricted to constructs printed in the FIB, as constructs printed in the cellulose-only bath could not be adequately removed after processing and the printed collagen filaments were no longer clearly distinguishable for reliable retrieval or downstream fibril analysis. Within FIB groups, following freeze-drying and construct retrieval, filament width was approximately 5% lower than immediately after printing across all groups (Figure 3c). Filament length after retrieval was close to the expected value of 25 mm, although the 1% AC-ECM groups printed at lower speeds showed filament lengths of approximately 21 mm (Figure 3d). No significant differences in filament length were observed between groups. Collagen fibril alignment increased with both ink concentration and print speed, with the highest value observed in the 5% AC-ECM group printed at 20 mm/s, reaching approximately 80% of fibrils aligning within ±10° of the printing direction compared with approximately 50% in the 1% AC-ECM group printed at the same speed (Figure 3e). SEM imaging showed collagen fibres with visible D-banding in both AC-ECM derived inks (Figure 3f).

**Figure 3:**
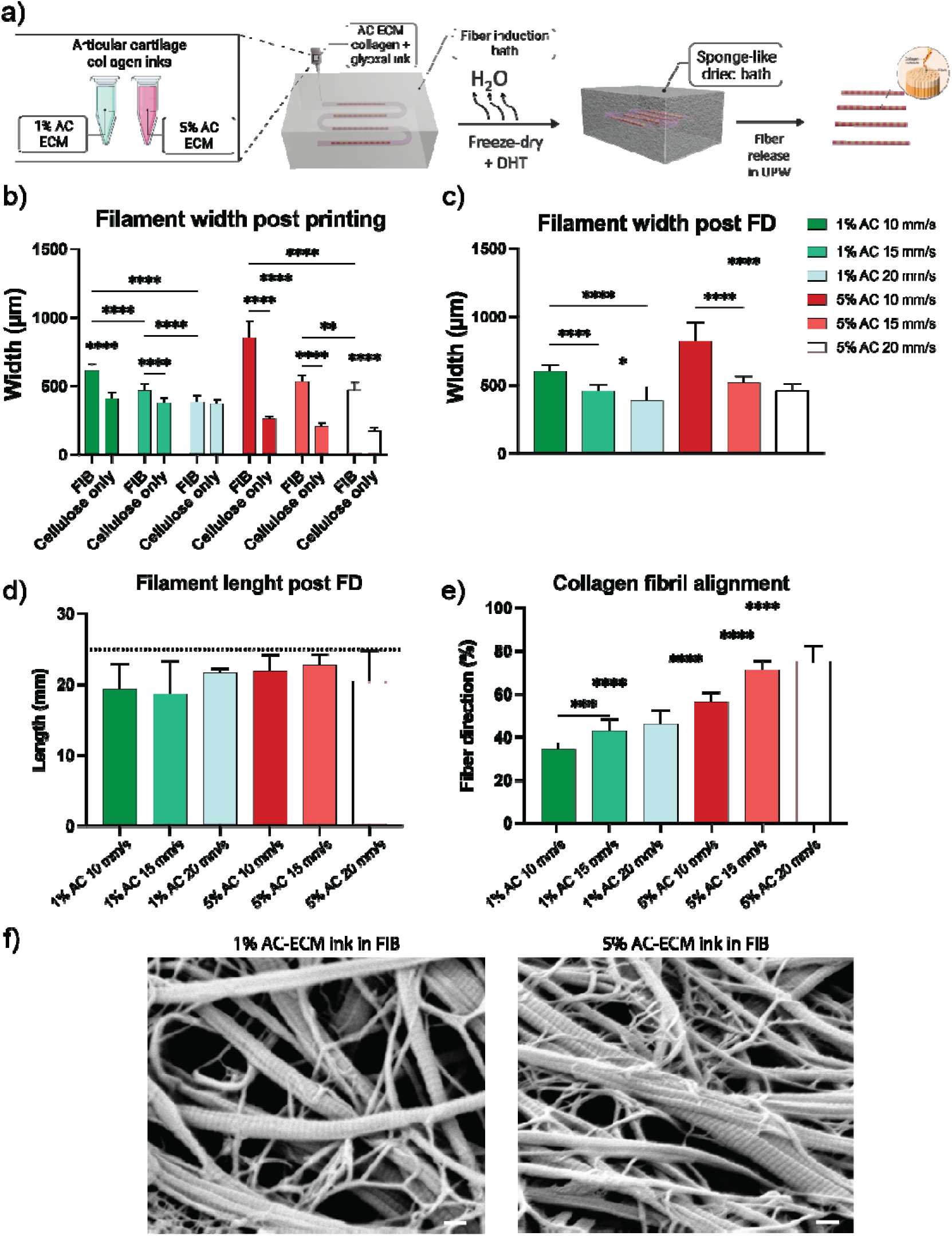
Embedded printing supports continuous filament deposition and directional fibrillar organisation of AC-ECM inks. Schematic of printing and post-processing workflow (a). Filament width immediately after printing in FIB and cellulose-only baths (b). Filament width after freeze-drying and construct retrieval from FIB bath (c). Filament length after construct retrieval from FIB bath (the dotted line indicates the expected filament length of 25 mm) (d). Percentage of collagen fibrils with local orientations lying within ±10° of the printing direction, quantified from SEM images using ImageJ directionality plugin (e). Representative SEM images of processed AC-ECM filaments showing collagen fibril morphology and D-banding (f). Data are presented as mean ± standard deviation, with statistical significance indicated in the figure. n = 3 printed samples per group. * p < 0.05, ** p < 0.01, *** p < 0.001, **** p < 0.0001. Scale bar = 200 nm.

### 3.3. Print parameters to control filament width, spacing and multilayer stacking fidelity

To assess the effect of print parameters on filament spacing and multilayer stacking, 1% and 5% AC-ECM inks were printed as single-layer or triple-layer line patterns at increasing filament spacing (Figure 4a). Representative images showed overlap, touching and separated filaments across the tested conditions (Figure 4b). Quantification showed that the spacing required to achieve overlap, touching and separation of the printed filaments was formulation-dependant (Figure 4c). For the 1% ink, filament separation was achieved at approximately 0.5 mm in single-layer prints but required a minimum spacing of approximately 0.75 mm in triple-layer prints. By contrast, the 5% ink showed no significant difference in the filament-to-filament spacing required to achieve separation between single-layer and triple-layer constructs.

**Figure 4:**
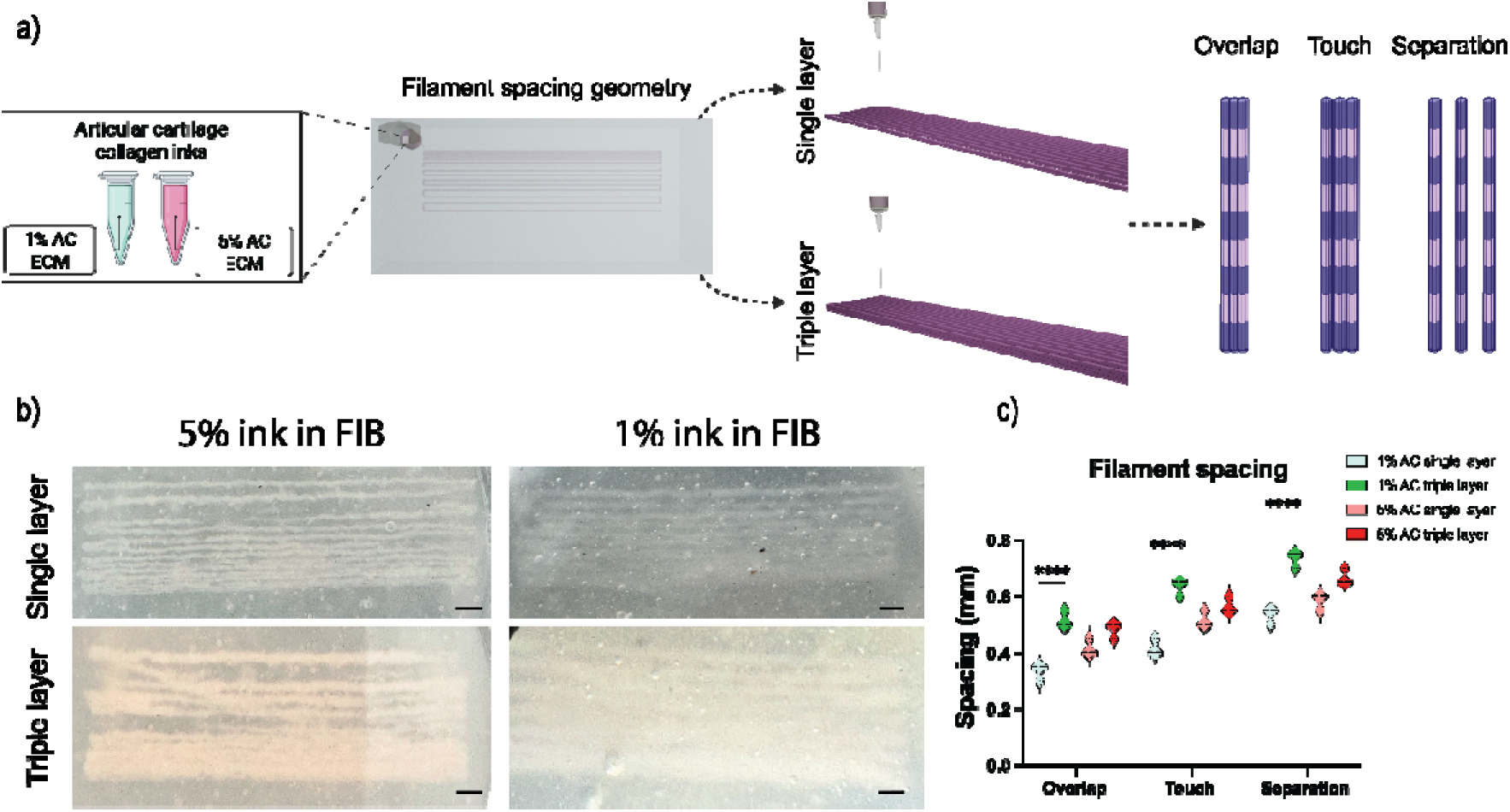
Print parameter effects on filament spacing and multilayer stacking fidelity. Schematic of the printing strategy used to generate adjacent filaments from 1% AC-ECM and 5% AC-ECM inks in single-layer and triple-layer formats (a). Representative images showing overlap, touching and separated filaments across the tested conditions (b). Quantified filament spacing required to achieve overlap, touching and separation for each ink formulation and layer number (c). Data are presented as mean ± standard deviation, with statistical significance indicated in the figure. n = 3 printed samples per group. **** p < 0.0001. Scale bar = 1 mm.

### 3.4. 3D printing of aligned sheets and scaffolds with programmed filament spacing

To assess whether the support bath could facilitate the 3D printing of larger structures, both 1% and 5% AC-ECM inks were used for printing cube-shaped scaffold constructs (Figure 5a). In addition, the 5% AC-ECM ink was used for printing 15mm by 15mm sheets (Figure 6a).

**Figure 5:**
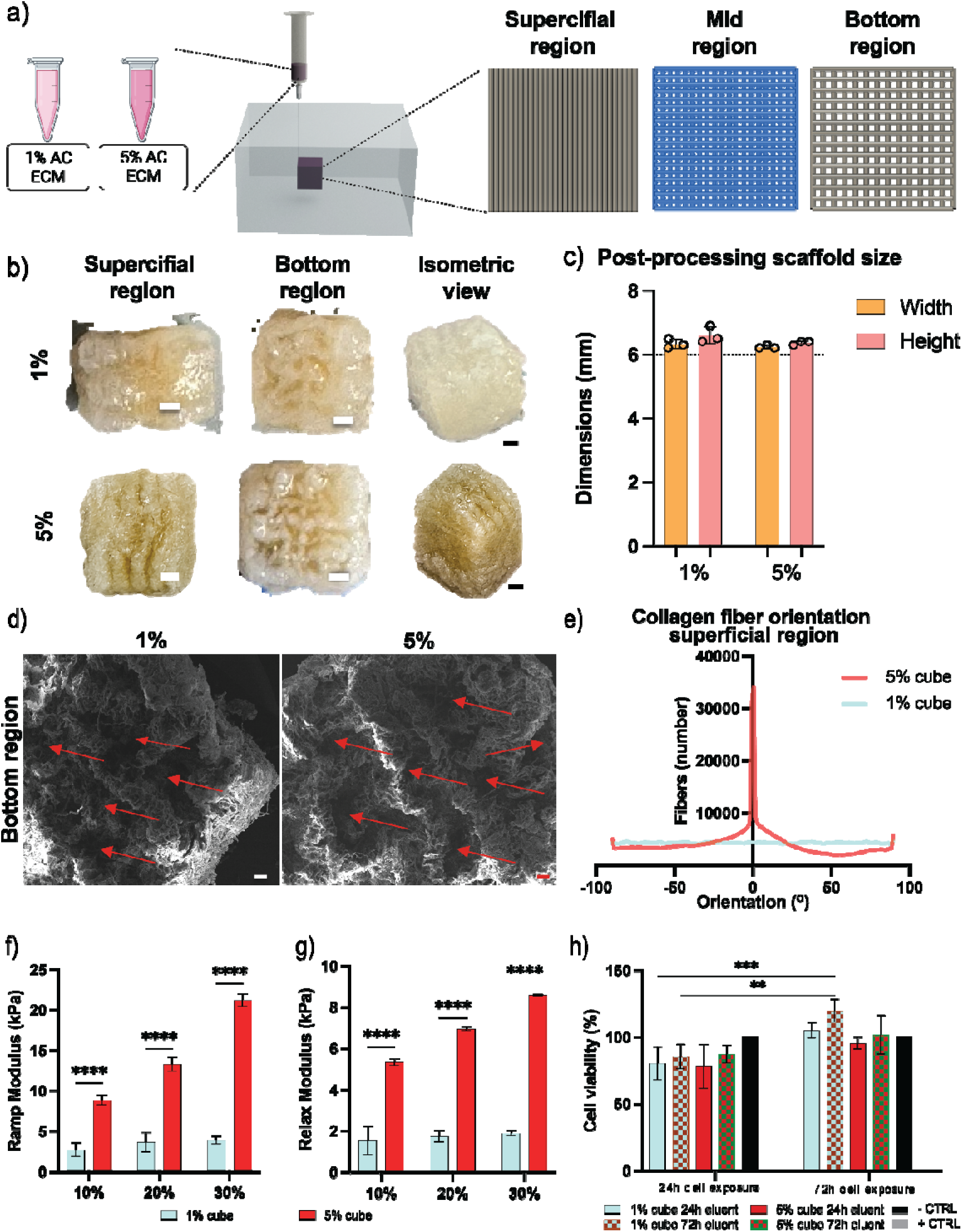
Construct-scale printing of a programmed spacing gradient AC-ECM scaffold with preserved overall dimensions and aligned collagen fibrils. Schematic illustration of the printing strategy used to fabricate the 6 × 6 × 6 mm scaffold with programmed variation in filament spacing through the construct height (a). Representative image of the retrieved scaffold and quantification of post-freeze-drying dimensions (b, c). Representative SEM image showing collagen fibrillar organisation within the scaffold. Red arrows point to pores within the scaffold (d). Fibre orientation analysis showing a dominant peak aligned with the printing direction (e). Mechanical properties of the retrieved constructs (f, g). Cell metabolic activity (%) after 24 h and 72 h of culture with construct extracts (h). n = 3 printed constructs per group. ** p < 0.01, *** p < 0.001, **** p < 0.0001. Scale bar = 1 mm (macro images), scale bar = 2 μm (SEM image).

**Figure 6:**
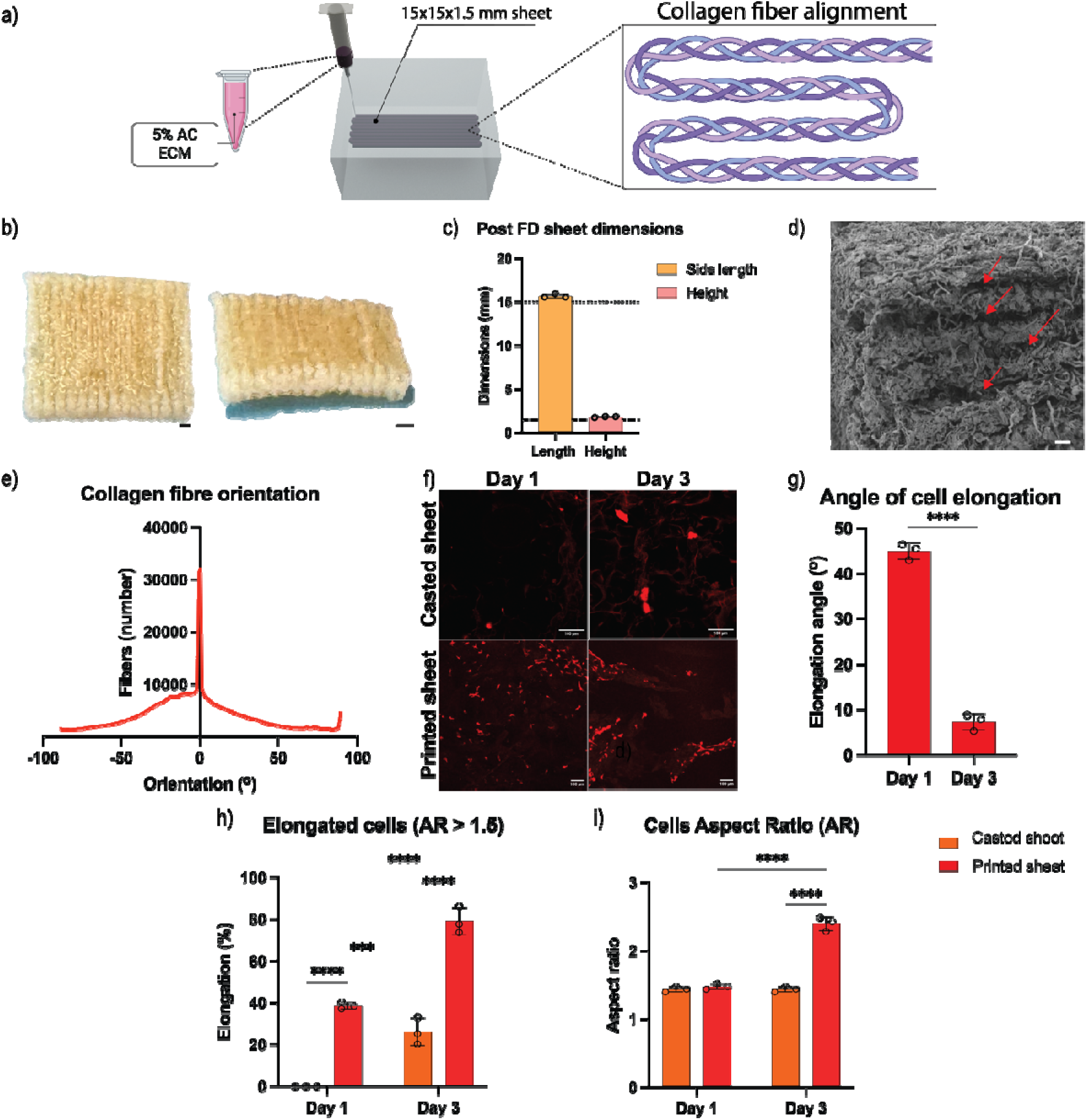
Construct-scale printing of a unidirectional AC-ECM sheet with aligned collagen fibrils. Schematic illustration of the printing strategy used to fabricate the sheet construct from 5% AC-ECM ink, showing parallel filament deposition and the intended filament architecture (a). Representative images of the retrieved construct and quantification of post-freeze-drying dimensions (b-c). Representative SEM image of the sheet construct showing collagen fibril organisation. Red arrows point to gaps between printed fibres (d). Fibre orientation analysis showing a dominant peak aligned with the printing direction (e). F-actin stain of MSCs seeded on casted and printed sheets at day 1 and day 3 (f). Angle of cell elongation, number of elongated cells and overall cells AR on the printed constructs (g-i). For construct dimensions, n = 3 printed constructs. *** p < 0.001, **** p < 0.0001. Scale bar = 1 mm (macro images), scale bar = 2 mm (SEM image), scale bar = 100 μm (F-actin).

The 6 × 6 × 6 mm scaffold was printed with a programmed variation in filament spacing through its depth (Figure 5a). After processing, the scaffold retained dimensions close to the intended design (Figure 5b, c). While the intended pore size gradient was not fully realised in the final construct, SEM imaging showed directional collagen fibrillar organisation within the scaffold (Figure 5d), which was supported by orientation analysis showing a dominant peak in the printing direction (Figure 5e). When compressed, the printed 5% AC-ECM constructs was significantly stiffer compared to the 1% AC-ECM constructs, with a ramp modulus and relaxation modulus up to fourfold higher than the lower concentration constructs (Figure 5f, g). Finally, MTT analysis showed that exposure to extracts collected from both formulations after 24 h and 72 h did not significantly reduce cellular metabolic activity relative to untreated controls, irrespective of the duration of cell exposure (Figure 5h).

Parallel filaments of the 5% AC-ECM ink were also printed into the support bath, thereby generating a continuous aligned structure (Figure 6a, b). Following freeze-drying and retrieval, the printed sheet retained dimensions close to the intended design, with a side length of approximately 15 mm and a height of approximately 1.5 mm (Figure 6b, c). Representative SEM imaging showed collagen fibrils with visible directional organisation (Figure 6d), while orientation analysis showed a dominant peak aligned with the printing direction (Figure 6e).

To assess if MSCs would respond to this aligned collagen structure, they were seeded onto the surface of the printed sheets. Over time in culture, MSCs became progressively more elongated and aligned parallel to the collagen filaments. At day 3, cells on printed patches exhibited a mean aspect ratio (AR) of approximately 2.5, significantly higher than that observed on the printed patches at day 1 and on the casted controls at day 3 (Figure 6i). Similarly, the proportion of elongated cells (AR > 1.5) increased from approximately 40% at day 1 to almost 80% at day 3 on the printed patches (Figure 6h). At day 3, the proportion of cells that was elongated was also significantly higher than that observed on the cast controls (∼30%). Finally, the absolute angular deviation between the cellular major axis and the collagen-filament direction decreased significantly from ∼45° at day 1 to ∼9° at day 3 (Figure 6g).

## 4. Discussion and conclusion

This study established a fibril-inducing support-bath (FIB) for embedded printing of collagen-rich AC-ECM inks, supporting filament deposition, post-processing, construct retrieval and biologically relevant anisotropy. Collagen-rich ECM inks often deform in air before stable fibrils develop because of low yield stress [33,69]. Embedded printing instead deposits them within a medium that yields around the nozzle and recovers afterwards [32]. Unlike passive support-baths, the FIB was designed to provide fibrillogenesis-promoting conditions [45]. Its contribution is therefore the integration of printing, fibril formation and stabilisation within one processing environment.

The NFC-based bath exhibited shear-thinning, shear recovery and a transition from solid-like to liquid-like behaviour at increasing strain. This agrees with the requirements for embedded printing, where the bath must yield locally and recover rapidly enough to prevent filament displacement or collapse [41,70]. Greater optical clarity than the MFC bath also enabled monitoring of nozzle position and deposition errors [69]. Selection of this formulation therefore balanced mechanical support with optical accessibility.

Printing remained concentration-dependent: the greater viscosity and viscoelastic moduli of the 5% AC-ECM ink likely reduced deformation and improved multilayer stability relative to the lower concertation ink. However, filament morphology was governed by the interaction between ink and bath rather than ink rheology alone. Filaments in FIB were wider than those in the cellulose-only bath, particularly for the 5% ink, however this was less evident for the 1% ink. This suggests that the FIB was not passive and that its ionic or macromolecular environment altered the ink after extrusion. Early fibrillar assembly, altered hydration, interfacial exchange or post-extrusion relaxation could all contribute to this observation [69,71]. These findings suggest that final print dimensions in such systems cannot be predicted without accounting for ink–bath interactions.

This dependence was reinforced by filament-spacing experiments, where the separation required to produce overlapping, touching or distinct filaments varied with ink concentration and layer number. This concentration-dependent stacking agrees with evidence that fidelity reflects ink rheology, extrusion, bath support and relaxation collectively [26,28]. CAD dimensions must therefore be calibrated against the processed construct. FIB also facilitated post-processing. Constructs in the cellulose-only bath could not be separated adequately after freeze-drying and DHT, whereas those in FIB could be stabilised while embedded and retrieved by immersion and washing. This avoided manipulation after rehydration and differs from FRESH, where gelatin microparticles are generally removed by warming [33]. Because gelatin can undergo modification during thermal dehydration [72], FIB may be more compatible with workflows requiring DHT before bath removal.

Large scale scaffolds could be printed using both the 1% and 5% AC-ECM inks, confirming that FIB accommodated inks with substantially different rheological properties. However the 5% AC-ECM ink supported noticeably greater collagen fibre alignment in the scaled-up scaffolds (Figure 5e). While overall print dimensions were retained, the programmed pore-size gradient was not clearly resolved. This separation between macroscopic and internal fidelity agrees with embedded-printing studies showing that microscopic porosity remains sensitive to printing speed, ink viscosity and the rheological relationship between ink and bath, even when the external geometry is preserved [71]. Filament spreading, layer fusion and dimensional changes during freeze-drying or DHT may have obscured the programmed spacing differences. This limitation is functionally relevant for the development of biomaterial scaffolds as pore architecture regulates permeability, nutrient transport, cell attachment and migration [73,74]. Greater separation between pore regions, CAD compensation for spreading and optimisation of the ink-to-bath viscosity ratio would likely lead to improvements in internal print quality.

SEM and orientation analysis demonstrated fibrillar organisation along the printing direction in both geometries. This agrees with evidence that shear and extensional forces during extrusion can orient the collagen-rich dECM along the deposition path [20], while ink rheology and support-bath interactions determine how effectively this organisation is retained [71]. Extrusion may therefore have initiated alignment, while FIB confinement limited relaxation and supported subsequent collagen fibrillation, with DHT stabilising the structure. This is notable because improving collagen-ink viscosity can enhance shape retention while restricting the molecular reorganisation required for alignment, and some approaches consequently require magnetic particles and external fields to generate anisotropy [78]. Here, alignment followed the programmed printing direction without an additional alignment step. Visible D-banding further supported preservation of collagen fibrillar assembly, structures which have previously been shown to provide directional cellular guidance [79].

Neither 1% nor 5% construct extracts significantly reduced cellular metabolic activity relative to untreated controls. This indicates no detectable short-term cytotoxicity from residual bath components, ECM-derived products or processing-associated leachables. Extract testing is consistent with established assessment of soluble substances released from biomaterials [80].

The observation that MSC aligned along the 3D printed sheets confirmed that the printed architecture provided topographical cues to the seeded cells. This agrees with literature showing that anisotropic collagen fibrils and filament-scale topography promote cell elongation through contact guidance, focal-adhesion organisation and cytoskeletal extension [78,81,82]. The reduction in absolute angular deviation from approximately 45° to 9° is particularly important because it demonstrates that cells did not simply elongate over time, but oriented specifically along the printed filaments. At day 3, MSCs on cast controls retained a lower aspect ratio, with significantly less cells elongating despite having the same AC-ECM composition. The observed cellular response is therefore more plausibly attributed to printing-induced anisotropy than material composition or culture duration alone. Longer term studies are required to access how the printed ECM sheets influence MSC differentiation[73].

A number of study limitations should also be highlighted. The mechanism causing increased filament width within the support bath was not resolved, the desired pore gradient was not preserved in the scaled-up scaffolds, and the SEM used to characterise collagen architecture in the printed constructs represents the dehydrated surface architecture. MTT assessed only soluble leachables, while cell culture evaluated only short-term surface responses. Future studies should examine hydrated and cyclic mechanics, internal fibril organisation, three-dimensional cell infiltration and longer-term cellular phenotype.

Overall, FIB combined rheological support, optical accessibility, collagen-compatible processing and construct retrieval. It accommodated a range of ink concentrations, preserved printing-induced fibrillar anisotropy, produced concentration-dependent mechanics without detectable extract cytotoxicity and importantly generated directional cues that guided MSC organisation. FIB therefore provides a platform in which embedded printing and collagen fibril are coupled to produce retrievable, anisotropic ECM constructs without the need for a separate alignment mechanism.

## ASSOCIATED CONTENT

### Authors contribution

Giovanni Gonnella: Conceptualization, Methodology, Investigation, Formal analysis, Writing.

Rosario Milazzo: Methodology, Formal analysis Rory Gibney: Methodology

Daniel J. Kelly: Conceptualization, Supervision, Writing. Review and editing, Funding acquisition.

### Declaration of generative AI and AI-assisted technologies in the manuscript preparation process

During the preparation of this work the author(s) used ChatGPT (OpenAI) in order to improve language and readability. After using this tool, the author(s) reviewed and edited the content as needed and take(s) full responsibility for the content of the published article.

### Data Availability Statement

All relevant data are within the manuscript and the Supporting Information. The data are available from the corresponding authors on reasonable request.

## Acknowledgments

Schematic diagrams of graphical abstract, figures 1-5 were created with BioRender.com, Autodesk Fusion360 software, Blender software, and Autodesk Illustrator software by G.G. This publication has emanated from research supported in part [or in whole] by a grant from Research Ireland under Grant number 12/RC/2278_P2. This project has received funding from the European Research Council (ERC) under the European Union’s Horizon Europe research and innovation programme (4D-BOUNDARIES; 101019344).

